# Integrative phylogenomic and pangenome landscape of *Bacillus*: insights from 10,000 genomes into taxonomy, functional potential, and biotechnological applications

**DOI:** 10.1101/2025.10.22.683932

**Authors:** Hector J. Acho-Vasquez, Sarah Henaut-Jacobs, Thiago M. Venancio

## Abstract

The genus *Bacillus* comprises Gram-positive, endospore-forming rods with a ubiquitous distribution across terrestrial, aquatic, and aerial environments, as well as associations with plants, animals, and food. To explore its diversity, we conducted an integrated phylogenomic and pangenome analysis using 10,839 publicly available RefSeq genomes (67 type strain genomes). Taxonomic delimitation combining genomic-distance metrics (Mash/ANI), network analyses, and a label-propagation algorithm assigned 10,276 genomes to operational communities, revealing novel complexes within *B. cereus* sensu lato and other clades. A robust phylogeny reconstructed from 103 representative genomes corroborated the ANI-based groupings. Forty-eight communities (≥10 genomes) were further analyzed for pangenome openness, showing a strong negative correlation between the saturation coefficient (α) and genomic fluidity (φ) (ρ = −0.636), indicating that open pangenomes exhibit high gene-content variability. Functional profiling revealed 135 antifungal genes and 15 secondary metabolite clusters, highlighting *B. velezensis*, *B. amyloliquefaciens*, and *B. subtilis* as rich reservoirs of hydrolytic enzymes, NRPS/PKS systems, and nutrient-competition traits. Additionally, 135 resistance determinants and 115 virulence factors were identified, mainly within *B. cereus* sensu lato. Biofertilization genes related to phosphorus, nitrogen, siderophore, potassium, and sulfur metabolism were broadly conserved, underscoring the genus biotechnological potential.

## INTRODUCTION

The genus *Bacillus* comprises Gram-positive, rod-shaped bacteria belonging to the family Bacillaceae (order Bacillales, class Bacilli, and phylum Firmicutes). These microorganisms are distinguished by their ability to form endospores and by their high metabolic plasticity, which enables growth under both aerobic and facultatively anaerobic conditions (Borriss, 2020; Göker & Oren, 2024; Maughan & Van der Auwera, 2011; Vos et al., 2009). The genus was first described by Cohn (1872) and currently includes more than 656 entries in the List of Prokaryotic names with Standing in Nomenclature Permanence (LPSN), of which 114 species have validly published names (Parte et al., 2020) (accessed July 2025).

*Bacillus* species are ubiquitously distributed and have been isolated from diverse habitats, including freshwater and marine environments, soils of various characteristics, air, animals, plants, and even food (Dérozier et al., 2023). Among the best characterized members are *B. cereus*, the etiological agent of foodborne illnesses such as emetic and diarrheal syndromes (Griffiths & Schraft, 2017; Yang et al., 2023); *B. anthracis*, the causative agent of anthrax (Bower et al., 2022); and *B. thuringiensis*, extensively used as a bioinsecticide in agricultural pest management (Sanahuja et al., 2011). These species belong to the *B. cereus* sensu lato group (Carroll et al., 2022). In contrast, members of the *B. subtilis* group (e.g., *B. subtilis* and *B. stercoris*) and the *B. amyloliquefaciens* group (e.g., *B. velezensis* and *B. amyloliquefaciens*) (Ngalimat et al., 2021) are recognized for their agricultural applications as plant growth-promoting bacteria (PGPB) and biocontrol agents with antifungal activity (Figueiredo et al., 2025; Li et al., 2023; Pengproh et al., 2023; Soliman et al., 2023; Teixeira et al., 2021). Ecological preferences within the genus appear to be shaped, at least in part, by genomic architecture, as organisms with similar lifestyles often share a higher proportion of genes (Henaut-Jacobs et al., 2023). However, accurately delineating these ecologically and functionally distinct groups remains challenging when relying solely on 16S rRNA gene-based taxonomy, whose limited resolution often obscures fine-scale relationships and misrepresents closely related taxa such as those within the *B. cereus* sensu lato complex (Chung et al., 2024; Maughan & Van der Auwera, 2011; Poretsky et al., 2014).

The rapid expansion of genome sequences in public repositories has enabled the adoption of whole-genome approaches such as average nucleotide identity (ANI), genomic distance metrics, and pan-genome analysis—methods that offer superior taxonomic resolution and allow inference of key aspects of strain lifestyle (Passarelli-Araujo et al., 2022, 2025). Recent studies demonstrate that such genome-based approaches are effective not only in identifying novel bacterial species but also for elucidating functional traits underlying ecological adaptation, as well as in characterizing genes associated with biocontrol, virulence, antimicrobial resistance, and plant growth-promoting traits (PGPTs). In this study, we analyzed 10,839 genomes, of which 10,276 high-quality genomes clusteredwere resolved into 103 distinct communities. We subsequently investigated the occurrence of 135 genes related to enzymes and secondary metabolites and 115 genes associated with virulence. The identification of these genetic determinants underscores the need for a rigorous, genome-informed selection of strains, paving the way toward the development of safe and sustainable agricultural bioinputs

## METHODS

### Dataset collection

A total of 10,839 publicly available *Bacillus* genomes from the NCBI RefSeq database in October 2024 using Datasets v.16.25.0 (https://www.ncbi.nlm.nih.gov/datasets/). Genome quality was assessed with CheckM v.1.2.3 (Parks et al., 2015), applying a minimum completeness threshold of 90% and a maximum contamination of 5%. BUSCO v5.8.0 (Simão et al., 2015) was additionally used with the Bacillales dataset as a reference, discarding genomes with completeness ≤90% or duplication ≥5 %.

### Type strain validation and network analysis

Type strains of the genus *Bacillus* were retrieved from the LPSN (Parte et al., 2020) to validate species names and to obtain the corresponding Genomic Taxonomy Database Designated Representative Genome (GTDB) representative genomes (Parks et al., 2018). Genomic distances for 1000-, 3000-, and 5000-bp sketches were computed using Mash v2.3.0 (Ondov et al., 2016). Reciprocal Mash distances were converted into genomic identity values (1 - Mash) and genome pairs with identities <0.95 were excluded, as they did not clearly resolve community structure.

Communities were identified using the label propagation algorithm (LPA) (Raghavan et al., 2007), as previously described (Passarelli-Araujo et al., 2022). Communities containing type genomes were named according to their corresponding validated type species (67 in total), whereas those lacking type genomes are assigned arbitrary labels (e.g. *Bacillus* sppN, where N corresponds to the number assigned to the community) (Henaut-Jacobs et al., 2023). In such cases, a representative genome was selected according to GTDB reference genome designation. To resolve highly connected or complex clusters, fastANI v1.34 (Jain et al., 2018) was applied for each complex separately. The resulting communities were compared with NCBI RefSeq database and GTDB classifications to identify potential misclassifications, which were subsequently inspected manually. For type strain comparisons, ANI was computed using pyANI v0.3.0.0-alpha using MUMmer-based alignment (Pritchard et al., 2016), ensuring that all genome pairs exhibited ≥95% identity. Community networks were visualized with the igraph v.2.1.4 package (Han et al., 2010).

### Phylogenetic reconstruction and pangenome analysis

All *Bacillus* genomes were re-annotated using Prokka v.1.14.6 (Seemann, 2014). Phylogenetic inference was performed using representative genomes from each community. Orthologous genes were identified with Orthofinder v. 3.0.1 (Emms & Kelly, 2015), and their sequences were aligned with MAFFT v7.526 (Rozewicki et al., 2019). The resulting alignments were concatenated and filtered using BMGE v1.12.1 (Criscuolo & Gribaldo, 2010). Phylogenetic inference was performed with IQ-TREE v.2.4.0 (Nguyen et al., 2015), employing the *TEST* option for automatic model selection. Node support values were computed from 1,000 nonparametric bootstrap replicates. *Metabacillus fastidiosus*, *M. litoralis*, and *M. niabensis* (accession number: GCF_001591625.1, GCF_007994985.1, GCF_030813165.1, respectively) were used as the outgroup. Phylogenies were visualized using iTOL v7 (Letunic & Bork, 2024).

Pangenome analysis was carried out for communities containing at least 10 genomes using Panaroo v.1.5.2 (Tonkin-Hill et al., 2020). Genomic openness was estimated by fitting a power law (α) model, and pangenome accumulation curves were generated with pagoo v0.3.18 (Ferrés & Iraola, 2021). Genomic fluidity, representing pairwise gene content diversity, was calculated with micropan v2 (Snipen & Liland, 2015).

### Identification and classification of genes involved in antifungal activity

A comprehensive literature review was performed to identify genes with experimentally validated antifungal activity in *Bacillus* strains. The corresponding coding sequences were retrieved from UniProt (Bateman et al., 2021) and NCBI Protein database (https://www.ncbi.nlm.nih.gov/protein/). Genes sharing the same name across different species were renamed by appending the initials of the respective species at the end (e.g., *chiA-Bv* for *B. velezensis*). When multiple genes with the same name were present in one species, a consecutive number was added (e.g., *chiA-Bv1*). Gene presence was determined with USEARCH v11.0.667 (Edgar, 2010), with thresholds of ≥60% identity and ≥50% coverage. Presence-absence matrices were generated and annotated in R with the tidyverse v.2.0.0 package (Wickham et al., 2019). To further evaluate the potential of these strains to produce antifungal secondary metabolites, antiSMASH v8.0.2 (Blin et al., 2025) was employed, considering similarity confidence values ≥50% (classified as Medium, 50–75%, and High, 75–100%).

### Identification of genes for antibiotic resistance, virulence, and biofertilization

Genes linked to antibiotic resistance, virulence, and biofertilization were identified using three specialized databases: the Comprehensive Antibiotic Resistance Database (CARD) (Alcock et al., 2023) for resistance genes, the Virulence Factor Database (VFDB) (B. Liu et al., 2022) for virulence factors, and PlaBAse (Patz et al., 2021) for biofertilization-related genes. All databases were downloaded on June 6, 2025. Gene detection was carried out with USEARCH v11.0.667 (Edgar, 2010) using thresholds of ≥60% identity and ≥50% coverage. Presence–absence matrices for all *Bacillus* genomes were generated in R with the tidyverse v2.0.0 package (Wickham et al., 2019).

## RESULTS

### Genome curation and taxonomic delimitation

We retrieved 10,839 *Bacillus* genomes from the NCBI RefSeq database (October 2024; Table S1). After applying stringent quality criteria (completeness ≥ 90% and contamination/duplication ≤ 5%), 10,625 genomes were retained, ranging from 2.7 to 8.0 Mb in size.

Sixty-seven type strains with validly published names, as listed in LSPN (April 2025), were identified, each having a corresponding reference genome in GTDB. For *B. paramobilis*, which lacks GTDB species assignment, a representative genome was manually selected (Table S2). Classification with LPA assigned 10,276 genomes into 78 *Bacillus* communities (Table S3, see Methods for details). Genomes excluded at this stage primarily belonged to closely related genera within Bacillaceae, such as *Niallia, Domibacillus, Pseudobacillus*, and *Priestia*. Each community was linked to a GTDB reference genome; those without a reference were labeled with the suffix (_XX).

Highly interconnected communities were grouped into species complexes, encompassing 76.5% of the dataset (Table 1). These complexes were further refined with FastANI (Table S4), yielding in 103 communities (34 within complexes). One representative genome from each community was subsequently analyzed with PyANI to achieve higher taxonomic resolution (≥ 95% identity threshold; Table S5). This analysis resolved nine communities as distinct species, while 25 remained as species complexes—three of which were unchanged after refinement (Figure 1).

**Figure 1.**
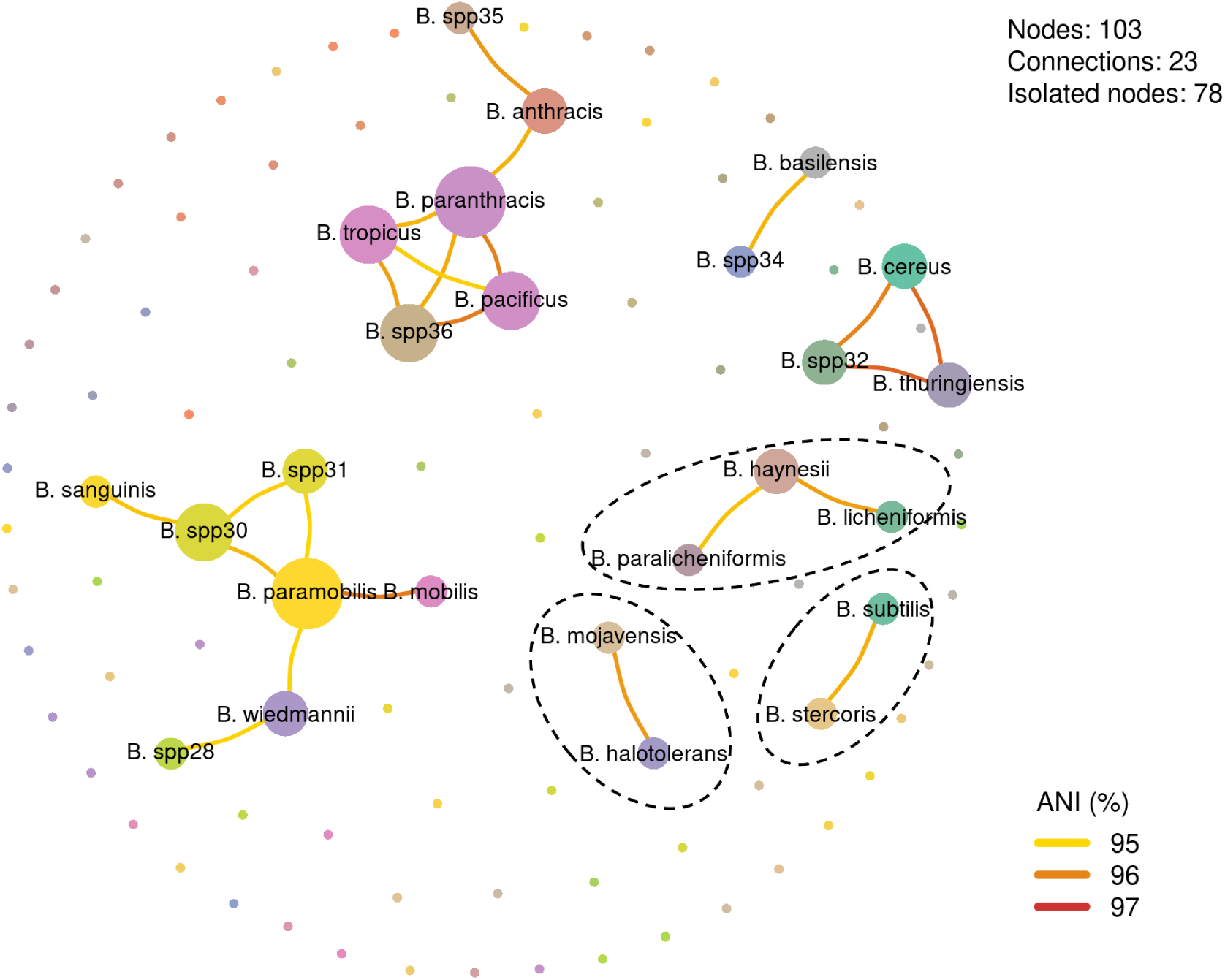
Network of genomic communities in the genus *Bacillus* based on average nucleotide identity (ANI). The network includes 103 representative genomes, of which 25 form 23 connections, while 78 remain isolated. Each node represents a genome, with node size proportional to its degree of centrality (number of connections). Edge colors denote ANI values: yellow (≥ 95%), orange (≥ 96%), and red (97%). Newly identified communities (e.g., *Bacillus* spp32, spp35, and spp36) show strong connectivity with reference species such as *B. cereus, B. thuringiensis, B. paranthracis*, and *B. paramobilis*, supporting their inclusion in the *B. cereus* sensu lato group or close phylogenetic proximity to recognized species (see also Figure 3). The communities delimited by dashed lines correspond to complexes that remained unchanged after refinement using pyANI. The remaining complexes were refined, including the resolution of 9 species (*i.e.*, separated communities)

**Table 1.** Species complexes in *Bacillus* resolved by community detection and ANI-based delimitation.

| Species complex | No. of genomes | ANI (%) |
| --- | --- | --- |
| *B. anthracis*, *B. basilensis*, *B. dicomae*, *B. pacificus*,<br>*B. paranthracis*, *B. sanguinis*, *B. tropicus*, *B. spp34*, *B.*<br>*spp35*, *B. spp36* | 1461 | 96.8 |
| *B. cereus*, *B. thuringiensis*, *B. spp32*, *B. spp33* | 2501 | 97.2 |
| *B. halotolerans*, *B. mojavensis* | 124 | 96.8 |
| *B. haynesii*, *B. licheniformis*, *B. paralicheniformis* | 776 | 95.8 |
| *B. hominis*, *B. mycoides* | 262 | 95.0 |
| *B. mobilis*, *B. paramobilis*, *B. wiedmannii*, *B. spp28*, *B.*<br>*spp29*, *B. spp30*, *B. spp31* | 416 | 96.2 |
| *B. nitratireducens*, *B. proteolyticus* | 101 | 95.0 |
| *B. siamensis*, *B. velezensis* | 1172 | 95.0 |
| *B. stercoris*, *B. subtilis* | 1055 | 96.8 |

Within the *B. anthracis* complex, the *B.* spp35 community was closely related to *B. anthracis* (ANI ≥ 95.77%), whereas *B.* spp36 exhibited multiple linkages to *B. paranthracis, B. tropicus*, and *B. pacificus* (ANI ≥ 95.53%), with *B. paranthracis* occupying a central, highly connected position. Likewise, *B. basilensis* and *B.* spp34 formed a distinct complex ANI ≥ 95.4%), clearly separated from the initial cluster. In the *B. cereus* complex (including *B. cereus* and *B. thuringiensis*), *B.* spp32 displayed strong connectivity with both species (ANI ≥ 96%).

In the *B. mobilis* complex, *B.* spp28 was linked to *B. wiedmannii* (ANI = 95.07%), while *B.* spp30 and *B.* spp31 were associated with to *B. paramobilis*, itself connected to *B. mobilis* and *B. wiedmannii* (ANI = 95.07%). Notably, *B.* spp30 is also connected to *B. sanguinis* (≥ 95.25%), which initially belonged to another complex (*B. anthracis*).

Conversely, the complexes formed by *B. haynesii*, *B. licheniformis*, and *B. paralicheniformis* (ANI ≥ 95.27%), as well as those involving *B. halotolerans* with *B. mojavensis* (ANI ≥ 95.75%) and *B. subtilis* with *B. stercoris* (ANI ≥ 95.47%), remained stable after PyANI refinement (Table 1).

Overall, the pyANI network analysis corroborated the Bacillus delineation obtained with FastANI while providing higher resolution within species complexes (Table 1, Table S4, Table S5).

### Species assignment correlation

To compare species assignments across classification frameworks, we analyzed genome counts derived from NCBI RefSeq, GTDB, and our graph-based LPA (Table S5). For species with validly published names listed in LPSN (as curated by GTDB, April 2025), Spearman’s rank correlation revealed strong and highly significant concordance among schemes. Genome counts assigned by RefSeq and GTDB were strongly correlated (ρ = 0.932, *p* < 2.35 × 10⁻^30^), as were those between RefSeq and LPA (ρ = 0.961, *p* < 7.64 × 10⁻^38^) and between GTDB and LPA (ρ = 0.969, *p* < 2.54 × 10⁻^41^) (Figure 2). These results indicate that, despite methodological differences, genome assignments are broadly consistent across reference-based (RefSeq, GTDB) and graph-based (LPA) taxonomic frameworks. Notably, LPA showed the highest concordance with GTDB, underscoring that the network-based approach effectively captures taxonomic relationships in agreement with genome-based species definitions.

**Figure 2.**
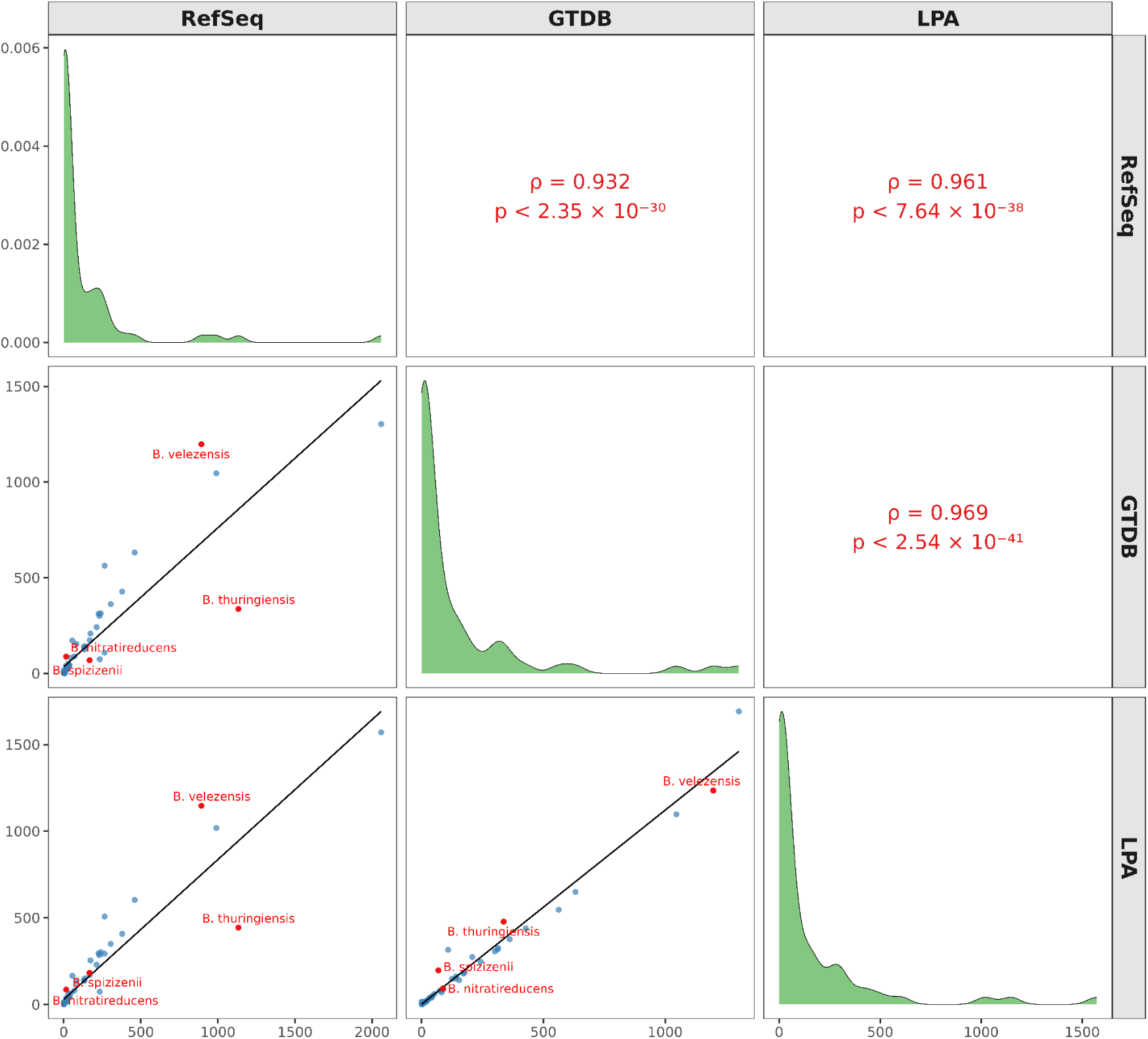
Taxonomic assignments among classification systems for selected *Bacillus* species. *B. thuringiensis* shows substantial overallocation in NCBI RefSeq (1,133 genomes) compared with GTDB (337) and LPA (444). *B. velezensis* is more inclusively defined by GTDB (1,198) and LPA (1,147) relative to RefSeq (893). In contrast, *B. spizizenii* is assigned significantly fewer genomes in GTDB (69) than in RefSeq (168) and LPA (183), suggesting a narrower taxonomic definition. *B. nitratireducens* appears underrepresented in RefSeq (16) compared to GTDB (88) and LPA (86), indicating potential misclassification or incomplete representation.

Nevertheless, marked discrepancies were observed for several taxa. For example, NCBI RefSeq classified 2,059 genomes as *B. cereus*, whereas GTDB and LPA assigned 1,303 and 1,571 genomes, respectively (Table S6, Figure 2). In addition *B. amyloliquefaciens* included 234 genomes in RefSeq, compared with only 74 in both GTDB and LPA. In contrast, *B. paranthracis* accounted for 265 genomes in RefSeq versus 563 in GTDB and 508 in LPA. Likewise, *B. subtilis* comprised 990 genomes in RefSeq compared with 1,045 in GTDB and 1,019 in LPA, whereas *B. velezensis* included 983 genomes in RefSeq versus 1,198 in GTDB and 1,147 in LPA. These discrepancies highlight persistent challenges in *Bacillus* species delimitation and reflect the divergent criteria applied across classification systems.

### Phylogenomic resolution of *Bacillus* communities

Phylogenetic reconstruction was conducted using one representative genome from each of the 103 communities (Table S2). ModelFinder, implemented in IQ-TREE, selected LG+F+I+G4 as the best-fit evolutionary model. Genomes of *Metabacillus* were used as outgroup taxa (see Methods). The resulting phylogeny showed strong branch support across all nodes (Figure 3). Species clustered according to their major operational groups, including the *B. cereus* sensu lato group (Carroll et al., 2022), *B. pumilus* (Fu et al., 2021), *B. amyloliquefaciens* (Ngalimat et al., 2021)*, B. licheniformis*, and *B. subtilis* (Caulier et al., 2019) groups. This resolution corroborates the network-based analysis (Figure 1), which also revealed complex communities. Overall, ANI-based species delimitation provided a robust framework for taxonomic resolution, while phylogenetic and functional group analyses offered complementary insights into species relationships and the biological basis of complex community formation. This integrative approach underscores the limitations of 16S rRNA-based classification, which lacks the resolution needed to discriminate among closely related *Bacillus* species (Liu et al., 2015).

**Figure 3.**
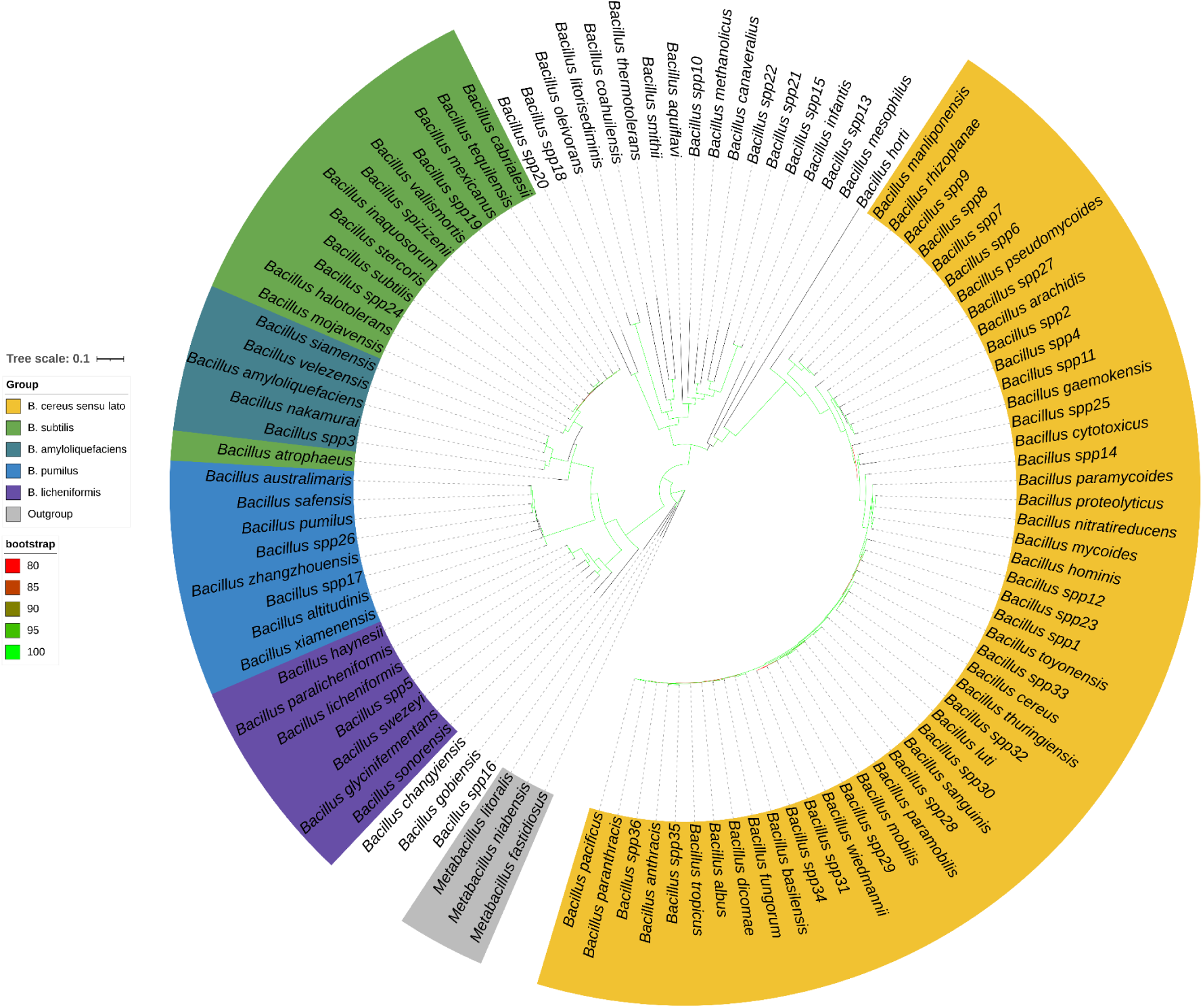
Maximum-likelihood phylogeny of 103 representative *Bacillus* genomes. Node support values ≥80% are indicated by green lines. Genome identifiers corresponding to each species are listed in Table S2. Major species groups are resolved and cluster according to their operational groups (e.g., *B. cereus* sensu lato, *B. pumilus*, *B. amyloliquefaciens*, *B. licheniformis*, and *B. subtilis*), in agreement with ANI-based community delimitation (see Figure 1).

### Pangenome architecture and genomic fluidity in *Bacillus*

Pangenome analysis was performed for 48 communities containing at least 10 genomes, enabling exploration of the relationship between genomic fluidity and the saturation coefficient (α) as indicators of genomic architecture and species dynamics. The pangenome, defined as the complete set of genes across all strains of a species or closely related group (Golchha et al., 2024; Rouli et al., 2015), is typically partitioned into three fractions: (i) the core genome, comprising genes present in all strains and linked to essential cellular functions and taxonomic identity; (ii) the accessory genome, consisting of genes shared by some but not all strains, often associated with niche adaptation, host interaction, or stress responses and resistance; and (iii) private or unique genes, found in a single strain, frequently reflecting specific adaptations or recent horizontal gene transfer (Golchha et al., 2024; Tettelin et al., 2008; Tettelin & Medini, 2020; Vernikos et al., 2015). Within this framework, genomic fluidity (φ) was calculated to quantify pairwise gene content dissimilarity, while α was used to assess pangenome openness. Lower α values denote open pangenomes with continuous gene acquisition, whereas higher values reflect closed, conserved pangenomes (Figure S1, Table S7). Spearman’s rank correlation revealed a significant negative association between α and genomic fluidity (ρ = −0.636; *p* = 1.19 × 10^⁻⁶^). A linear regression confirmed this trend (*F*₁.₄₆ = 41.28; *p* = 6.66 × 10⁻⁸), explaining 46.1% of the variance (adjusted R² = 0.461). The resulting equation, φ = 0.4499 − 0.4242α, suggests that genomic fluidity decreases by approximately 0.42 units for every unit increase in α (Figure 4).

**Figure 4.**
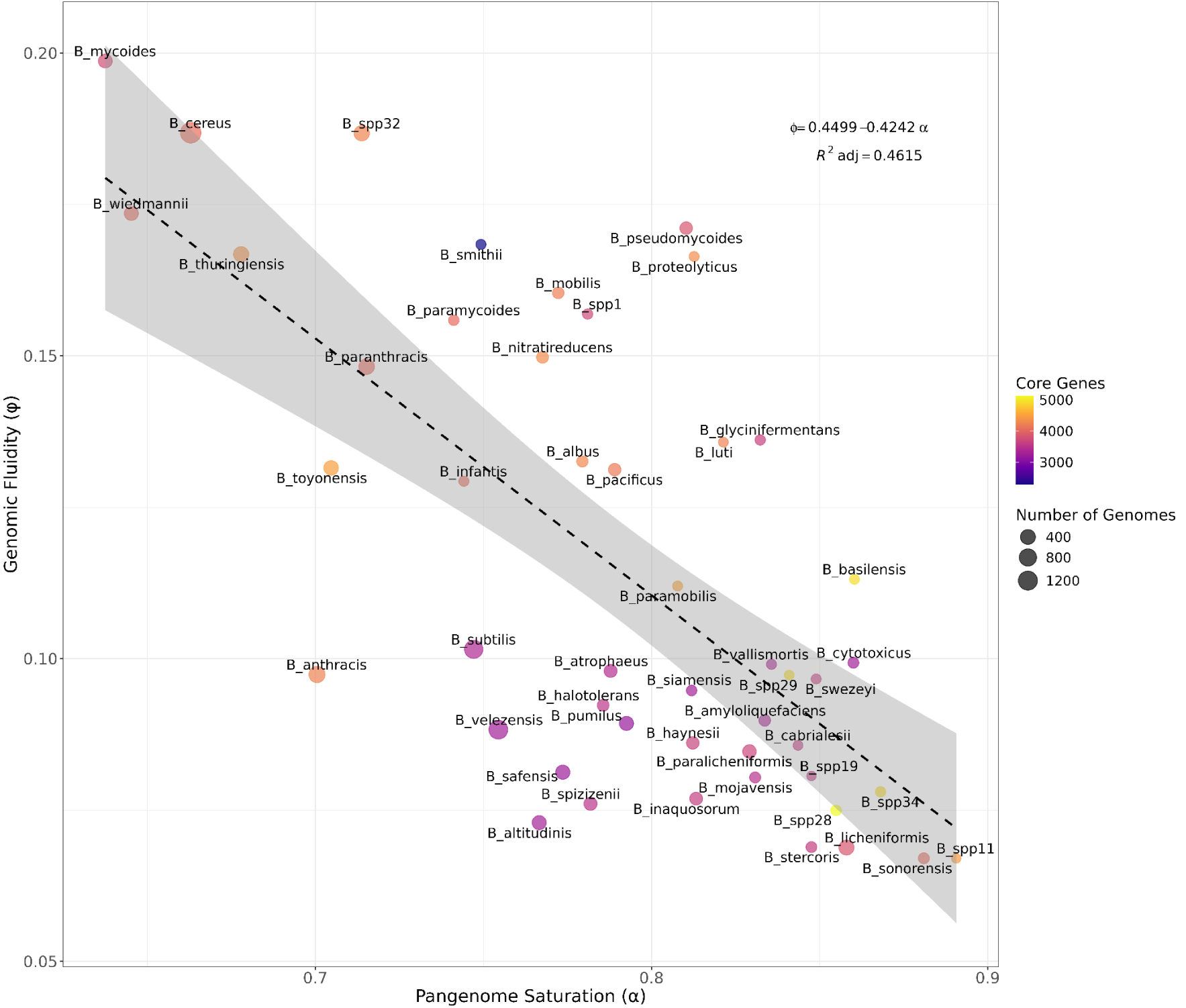
Pangenome openness and genomic fluidity in *Bacillus* species. (a) Cumulative pangenome growth curves of 48 *Bacillus* communities with at least 10 genomes, illustrating the rate of novel gene discovery as additional genomes are incorporated and highlighting variation in pangenome openness among species. (b) Relationship between pangenome openness (saturation curve coefficient, α) and genomic fluidity (φ). The linear regression model (φ = 0.4499 − 0.4242α) revealed a significant negative association (F₁,₄₆ = 41.28, p = 6.66 × 10⁻^8^), showing that species with more open pangenomes tend to exhibit higher genomic fluidity.

Interestingly, species such as *B.* spp11*, B. sonorensis, B.* spp34*, B. basilensis*, and *B. cytotoxicus*—all members of the *B. cereus* sensu lato group (Carroll et al., 2022)—displayed relatively high α values and a larger proportion of core genes, a pattern that may reflect the limited number of genomes analyzed and/or genuine ecological specialization. By contrast, *B. licheniformis* and *B. paralicheniformis* exhibited the most convincingly “closed” pangenomes, supported by extensive sampling (408 and 230, respectively) and consistently high α values. Conversely, species with low α values, including *B. mycoides* (n = 255), *B. cereus* (n = 1,592), *B. thuringiensis* (n = 444), and *B. wiedmannii* (n = 285), showed highly open pangenomes and elevated genomic fluidity (Table S6). This pattern is consistent with a generalist lifestyle, extensive genomic plasticity, and higher rates of HGT, enabling rapid adaptation to diverse environments. Notably, this group includes opportunistic pathogens (e.g., *B. cereus* group), where genetic versatility likely contributes to ecological success and pathogenic potential (Walker-York-Moore et al., 2017). Collectively, these results indicate that pangenome openness and genomic fluidity in *Bacillus* are shaped by a combination of sampling depth and species-specific ecological strategies.

### Secondary metabolites and competitive antifungal strategies in *Bacillus*

Across the 103 communities analysed (10,276 genomes in total), three major antifungal mechanisms were identified: (i) cell wall degradation, via hydrolytic enzymes, (ii) growth inhibition mediated by peptides and secondary metabolites, (iii) competitive inhibition through nutrient sequestration. In total, 135 genes involved in these mechanisms were evaluated (Figure S2, Tables S8, S9 and S10).

The first mechanism, cell wall degradation, targets fungal structural polysaccharides. Chitinases and glucanases play central roles in breaking down chitin and glucans, which are essential components of fungal cell walls (Ruiz-Herrera & Ortiz-Castellanos, 2019; Synowiecki & Al-Khateeb, 2003). Additionally, exposed chitin can be converted into chitosan, a pathogenicity strategy employed by fungi infecting both plants and humans (e.g. *Colletotrichum graminicola* and *Cryptococcus neoformans*) (Baker et al., 2011; El Gueddari et al., 2002; Nampally et al., 2012). Genes encoding chitinases (e.g., *chiA*, *chiB*, *chiD*), glucanases (e.g., *bglA, bglS*), and a chitosanase (e.g., csn), were widely distributed and often present as multiple homologs across *Bacillus* species. These enzymes act complementarily, degrading chitin, β-glucans, and chitosan. When evaluated individually, however, their activity is limited (Mauch et al., 1988), for instance, synergistic activity markedly enhances antifungal effects, as demonstrated in *B. subtilis* and *B. methylotropicus* (syn. *B. velezensis*), which suppress the growth of *Alternaria triticina* and *Bipolaris sorokiniana* through combined chitinase-glucanase production (Saini et al., 2024). Likewise, chitosanase in *B. subtilis* has been shown to inhibit *Botrytis cinerea, Fusarium oxysporum*, and *F. solani* (Pang et al., 2021; Park et al., 2008).

The highest potential for chitinase-mediated degradation was observed in the *B. cereus* sensu lato clade, with nine genes identified, particularly in *B. toyonensis, B. thuringiensis, B.* spp32*, B. cereus, B. wiedmannii*, and *B. tropicus*. In contrast, smaller chitinase repertoires (seven genes) were detected in the *B. pumilus, B. amyloliquefaciens, B. licheniformis*, and *B. subtilis* groups, where the presence of the *chiA-Bv* gene was notable (Tran et al., 2022). Interestingly, *B. paranthracis*, a member of the B. cereus sensu lato clade, also harbored this gene despite its divergence from more distant phylogenetic lineages. By comparison, *B. velezensis, B. siamensis*, and *B. amyloliquefaciens* (*B. amyloliquefaciens* group), together with *B. atrophaeus, B. mojavensis*, and *B. halotolerans* (*B. subtilis* group), exhibited greater enzymatic potential associated with chitosanases and glucanases. Notably*, B. atrophaeus, B. mojavensis*, *B. halotolerans, Bacillus* spp3*, B. velezensis, B. siamensis*, and *B. amyloliquefaciens* harbored at least two genes from each of the three hydrolytic categories, indicating broad antifungal potential.

The second antifungal mechanism involves the production of antifungal peptides and secondary metabolites. Non-ribosomal peptide synthetases (NRPSs) encode compounds such as surfactin (*srfA*), bacillomycin (*bmy*), fengycin (*fen*), iturin (*itu*), and plipastatin (*pps*), which bind to fungal membranes and disrupt their integrity (Cawoy et al., 2014; Jourdan et al., 2009; Khatoon et al., 2022). Polyketide synthase (PKS) clusters, responsible for difficidin (*dfn*) and macrolactin (*mln*) production, were also detected; these metabolites interfere with cell envelope biosynthesis and nucleic acid or protein synthesis (Khatoon et al., 2022; Kumariya et al., 2019; Papagianni, 2003). Hybrid NRPS/PKS pathways, including *pks* (bacillaene), *myc* (mycosubtilin), and *zma* (zwittermicin), combine polyketide and peptide modules to yield structurally complex compounds with dual activity against fungal membranes (Abdelmoteleb et al., 2017; Emmert et al., 2004; Kevany et al., 2009). Ribosomally synthesized and post-translationally modified peptides (RiPPs), such as subtilin (*spa*) and subtilosin (*alb* + *sbo*), were also identified; both act by inducing membrane permeabilization and apoptosis (Klein et al., 1992; Marx et al., 2001). In addition, rhizocticin (*rhi*), a phosphonopeptide that acts synergistically with the protease subtilisin (*aprE*), was observed. Together, they inhibit filamentous and budding fungi by interfering with intracellular metabolism (Abdelmoteleb et al., 2017; Borisova et al., 2010; J. Hu et al., 2022; Kugler et al., 1990).

Analysis with antiSMASH v8.0.2 confirmed the presence of 15 secondary metabolite clusters associated with antifungal activity (Figure 5, Table S11). Fengycin was the most widespread metabolite, detected in 33 species including *B. altitudinis*, *B. amyloliquefaciens*, and *B. atrophaeus*. Surfactin was present in 17 species, while mycosubtilin was restricted to five (*B. atrophaeus*, *B. inaquosorum*, *B. nakamurai*, *B. spizizenii*, and *B. velezensis*). Bacillomycin D occurred only in *B. mexicanus* and *B. velezensis*, whereas plipastatin was identified in nine species. Subtilosin A was found in 12 species, and zwittermicin A was limited to *B. cereus*, *B.* spp32, *B. thuringiensis*, and *B. wiedmannii*. Difficidin was exceptionally rare, occurring only in *B. siamensis* and *B. velezensis*. Macrolactin H and rhizocticin A were present in six species, while subtilin was identified in seven species, including *B. amyloliquefaciens* and *B. subtilis*. Notably, the Iturin cluster was not detected by antiSMASH due to its low similarity (<50%) to MIBiG reference clusters, which prevented its annotation as a high-confidence biosynthetic gene cluster. In contrast, the USEARCH analysis identified the complete cluster in species such as *B. amyloliquefaciens*, *B. siamensis*, and *B. velezensis* (Figure S2). This discrepancy arises from fundamental methodological differences: antiSMASH integrates multiple layers of information through ClusterCompare and KnownClusterBlast, including the presence of biosynthetic domains, NRPS/PKS module architecture, gene functions, sequence identity, and synteny to produce a composite similarity score. By contrast, USEARCH relies solely on direct protein-level alignments, reporting identities above 60% with coverage >50%. As a result, even when individual genes show strong similarity, the aggregate score calculated by antiSMASH may remain low, potentially underestimating the functional presence of this biosynthetic gene cluster.

**Figure 5.**
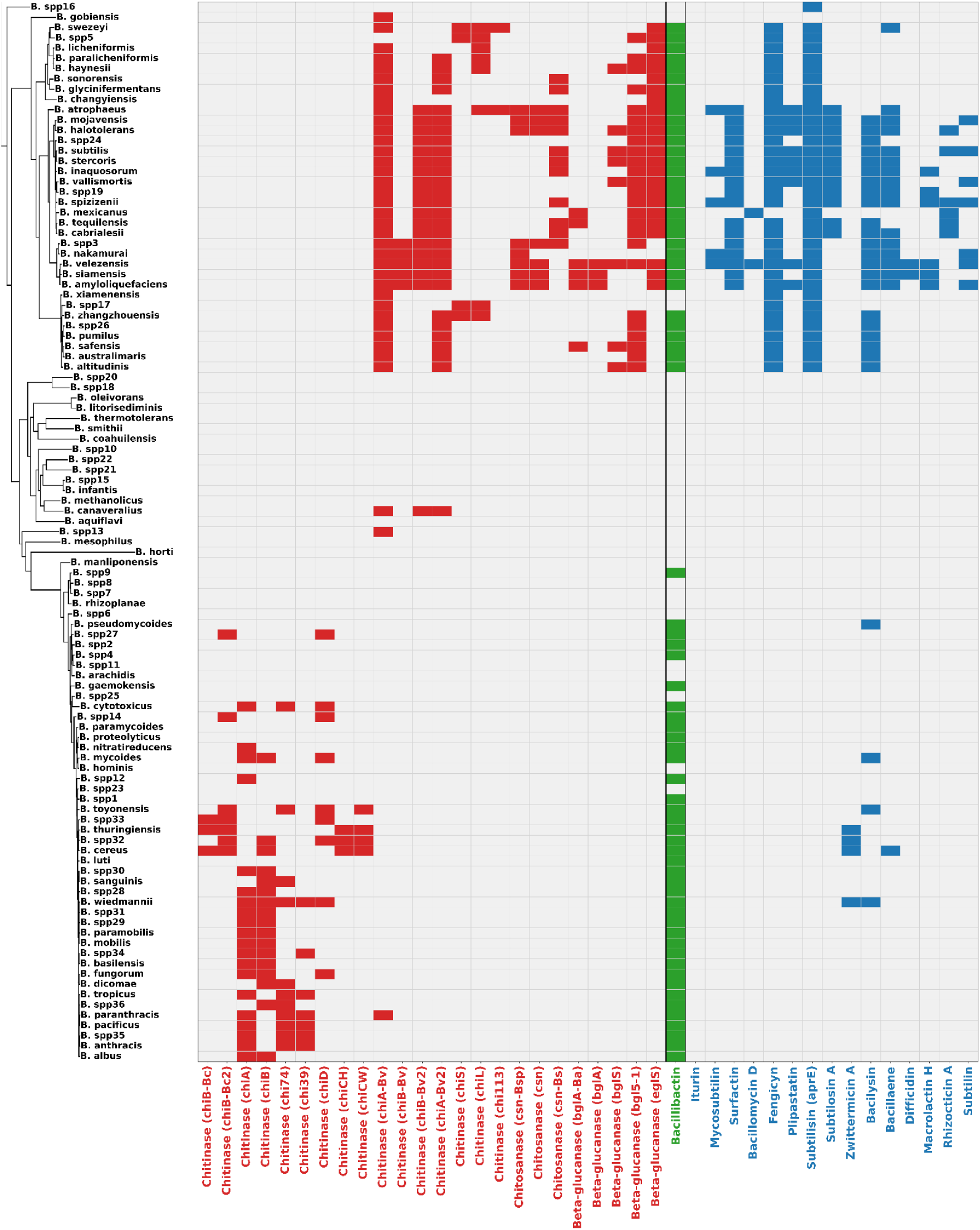
Presence–absence profiles of chitinases, glucanases, glucosidases, and antifungal peptide biosynthetic clusters across *Bacillus* species. Genes listed with locus identifiers (e.g., chitinase *chi74*, subtilisin *aprE*) were detected using USEARCH v11.0.667. Metabolite names without gene identifiers (e.g., bacillibactin, surfactin, fengicyn) correspond to biosynthetic clusters predicted with antiSMASH v8.0.2 (Table S9). Categories: Cell wall degradation (red), Competitive inhibition (green) and Growth inhibition (blue).

The third antifungal mechanism involved nutrient competition, such as Bacillibactin (*dhb*), a siderophore responsible for iron acquisition (Dimopoulou et al., 2021; Manetsberger et al., 2025), was present in 71 species, although partial operon losses (e.g., *dhbB*, *dhbC* by USEARCH) were occasionally observed. Thirty-two species lacked the operon entirely, including *B. aquiflavi*, *B. arachidis*, and *B. coahuilensis*. Together, these results show that antifungal potential is unevenly distributed across the genus. Notably, *B. velezensis, B. siamensis, B. amyloliquefaciens, B. spizizenii*, *B. subtilis*, and *B. atrophaeus* exhibited the broadest repertoire of antifungal genes, encompassing all three major mechanisms.

### Antibiotic resistance landscape across *Bacillus*

Analysis of antibiotic resistance determinants across 10,276 genomes identified 135 unique genes (Figure 6, Table S12). Genes involved in antibiotic inactivation were of particular clinical relevance. Among these, TEM β-lactamase variants (e.g., *TEM-1, TEM-116, TEM-181, TEM-229*) stood out, as they are widely reported in enterobacteria such as *Enterobacter, Citrobacter, Klebsiella*, and *Proteus*, where they act as major drivers of β-lactam resistance (Naidoo et al., 2020; Oduro-Mensah et al., 2016; Salverda et al., 2010). In addition, intrinsic β-lactamases characteristic of *Bacillus* were detected, including *BcII*, *Bla1*, and *Bla2*, were detected and are known to confer basal resistance to penicillin and ampicillin (Zheng et al., 2024). Collectively, these genes provide resistance to penicillins, cephalosporins, and carbapenems, and, in the case of TEM variants, also to monobactams. They were identified in several species, including *B. safensis, B. thuringiensis, B. altitudinis, B. cereus*, and *B. pacificus*. Additional inactivation determinants included FosBx1 and FosB, linked to fosfomycin resistance, which were exclusive to the *B. cereus* sensu lato group (Kowalska et al., 2024; Roberts et al., 2013).

**Figure 6.**
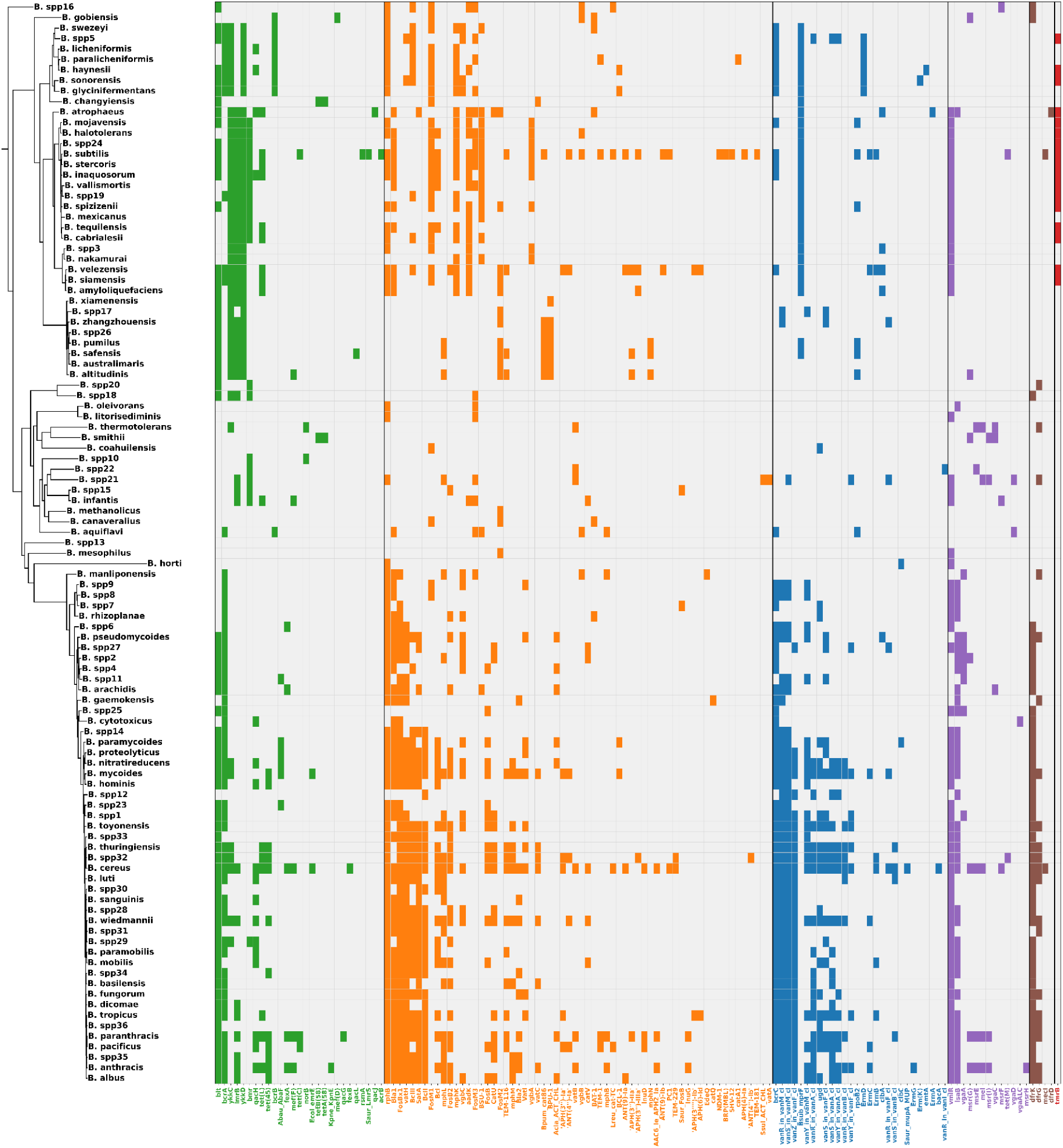
Distribution of antibiotic resistance mechanisms across *Bacillus* species based on CARD database annotations. Seven distinct resistance mechanisms were identified: antibiotic efflux (green, 27 genes), antibiotic inactivation (orange, 62 genes), antibiotic target alteration (blue, 28 genes), antibiotic target protection (purple, 13 genes), antibiotic target replacement (brown, 4 genes) and reduced permeability to antibiotic (red, 1 gene).

Genes associated with antibiotic target alteration were also observed, most notably the *vanA* and *vanB* operons and their regulatory components (*vanR*, *vanS*, *vanY*, *vanZ*), which mediate glycopeptide resistance by replacing the terminal D-Ala-D-Ala dipeptide with D-Ala-D-Lac, thereby reducing vancomycin binding affinity (Hill et al., 2010; Moosavian et al., 2018; McInnes et al., 2024). In our dataset, these operons were detected in species such as *B. albus, B. anthracis, B. mycoides, B. basilensis*, and *B. cereus*.

The antibiotic efflux mechanism was represented by tetracycline resistance genes, including *tet(M)*, *tet(L)*, *tet(45)*, *tetA(58)*, and *tetB(58)*, which were identified in members of the *B. cereus* group as well as in streptococci (Agersö et al., 2002; Hedayatianfard et al., 2014; X. Hu et al., 2022; Jeong & Lee, 2021). In the *B. amyloliquefaciens* group, *tet(L)* conferred only low-level resistance to tetracycline, as it was not associated with mobile genetic elements and is therefore considered an intrinsic determinant with very limited risk of horizontal transfer (Nøhr-Meldgaard et al., 2022). These genes were detected in multiple species, including *B. cereus, B. infantis, B. anthracis, B. cabrialesii, B. atrophaeus*, and *B. amyloliquefaciens*, highlighting the potential for widespread efflux-based resistance. Additional efflux-related determinants included *bcrA*, *bcrB* and *bcrC* belonging to Antibiotic target alteration mechanism, which form an ABC transporter system that provides self-protection against bacitracin by actively exporting the antibiotic. First characterized in *B. licheniformis*, a natural bacitracin producer (Neumüller et al., 2001; Podlesek et al., 1995), this system was also present in *B. paralicheniformis, B. haynesii*, and *B. sonorensis* (all members of the *B. licheniformis* group). Furthermore, *ykkC* and *ykkD,* genes implicated in chloramphenicol resistance in *B. subtilis* (Jin et al., 2025), were identified in the *B. subtilis, B. amyloliquefaciens*, and B. *pumilus* groups.

In addition to efflux systems, genes associated with antibiotic target alteration were also observed. Notably, *vanZ_in_vanF_cl* genes were identified, conferring resistance to glycopeptides such as vancomycin (Hill et al., 2010; Moosavian et al., 2018).

Finally, antibiotic inactivation mechanisms were represented by the *BcII* β-lactamase, which provides resistance to penicillins and cephalosporins (Zheng et al., 2024). Two additional genes, *FosBx1* and *FosB,* confer resistance to fosfomycin (Kowalska et al., 2024; Roberts et al., 2013) and were found exclusively in the *B. cereus* sensu lato group.

### Virulence gene repertoire and distribution within the genus *Bacillus*

Analysis of virulence-associated genes revealed 115 distinct loci distributed across the genus (Figure 7, Table S13). According to VFDB, these genes were classified into several functional categories, with exotoxins being the most prominent. Exotoxin genes were largely restricted to the *B. cereus* sensu lato group and included *alo*, which encodes a cholesterol-dependent cytolysin in *B. anthracis*. This toxin binds to host membranes, induces structural rearrangements, forms pores, and ultimately causes cytolysis (Mosser & Rest, 2006). Additional exotoxin genes included *hblA*, *hblC*, and *hblD*, which are related with hemolytic and cytotoxic activities (Ramm et al., 2021). Likewise, *nheA*, *nheB*, *nheC,* and *cytK*, encoding non-hemolytic enterotoxins and cytotoxins, were also detected. Together, these genes are associated with diarrheal and food poisoning caused by *B. cereus* (Lindbäck et al., 2004; Mostafa et al., 2024; Ramm et al., 2021). Exotoxin determinants were detected in multiple species, including *B. albus, B. anthracis, B. pacificus, B. paranthracis*, and *B. cereus*.

**Figure 7.**
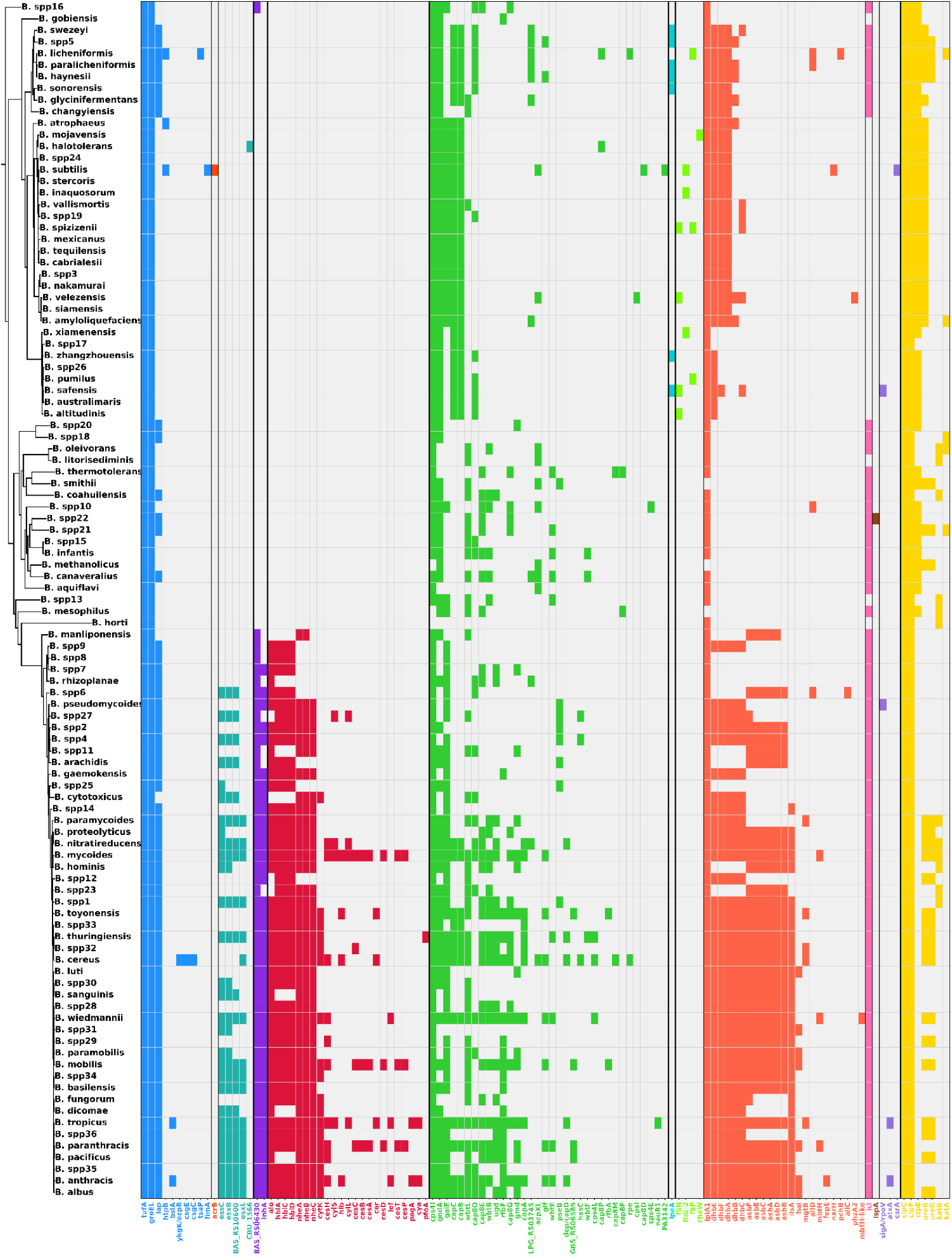
Functional categories of virulence factors identified across *Bacillus* species, based on VFDB annotations. A total of 13 categories were detected: Adherence (blue, 10 genes), Antimicrobial activity/Competitive advantage (dark orange, 1 gene), Effector delivery system (turquoise, 5 genes), Exoenzyme (violet, 2 genes), Exotoxin (red, 23 genes), Immune modulation (green, 33 genes), Invasion (light blue, 1 gene), Motility (lime green, 4 genes), Nutritional/Metabolic factor (coral, 23 genes), Others (fuchsia, 1 gene), Post-translational modification (brown, 1 gene), Regulation (lilac, 3 genes) and Stress survival (yellow, 7 genes). The Exotoxin and Exoenzyme categories were largely restricted to the *B. cereus* sensu lato group, reflecting its pathogenic potential.

Exoenzymes were represented by *inhA* and its homolog *BAS_RS06430*, encoding zinc-dependent metalloproteases with cytotoxic and exocytic activity. These proteins promote bacterial multiplication, immune evasion, and persistence, contributing to gastrointestinal and non-gastrointestinal infections caused by *B. cereus* (Guillemet et al., 2010; Ramarao & Lereclus, 2005). These genes were consistently present in *B. cereus* sensu lato species, including *B. cereus, B. albus, B. anthracis, B. sanguinis, B. paranthracis*, and *B. tropicus*.

The Effector Delivery System category was defined by the type VII secretion system (T7SS), which secretes small, signal peptide–independent proteins (∼100 amino acids). Key components include EssC, EsxB, and EsxL. EssC encodes a membrane-associated ATPase required for substrate recognition and secretion, while EsxB and EsxL represent WXG100 family proteins typically secreted as heterodimers. In Bacillus and related Firmicutes, T7SSs are primarily associated with interbacterial antagonism and niche competition, whereas in Mycobacterium tuberculosis, homologous systems contribute to immune evasion by suppressing CIITA/MHC-II expression and impairing macrophage activation (Bowran et al., 2023; Fan et al., 2015; Jäger et al., 2018; Sengupta et al., 2017). The T7SS was identified in species such as *B. paramycoides, B. nitratireducens, B. mycoides, B. hominis, B. wiedmannii*, and *B. basilensis*.

Finally, nutritional/metabolic virulence factors were represented by the *asb* operon (*asbA-F*), responsible for the biosynthesis of the siderophore petrobactin. Petrobactin contributes to iron acquisition, oxidative stress protection, and sporulation, thereby facilitating transmission in *B. anthracis* (Cendrowski et al., 2004; Hagan et al., 2018; Nusca et al., 2012). In addition to *B. anthracis*, we detected the operon in *B. toyonensis, B. thuringiensis, B. cereus, B. sanguinis, B. wiedmannii,* and *B. paramobilis*. The *ilsA* gene in *B. cereus* encodes a cell-surface protein containing NEAT (Near-Iron Transporter), LRR (Leucine-Rich Repeat), and SLH (Surface Layer Homology) domains, which facilitate iron acquisition under iron-limited conditions during host infection (Abi-Khalil et al., 2015; Koehler, 2009).

Collectively, these findings show that virulence-associated genes are unevenly distributed across *Bacillus*, with the majority concentrated in the *B. cereus* sensu lato group, where they likely underpin both pathogenic potential and ecological success.

### Genetic basis of biofertilization potential in *Bacillus*

Genes associated with plant growth-promoting traits were analyzed using the PLaBAse database, with a focus on biofertilization. Figure 8 and Table S14 summarize the detected genes, which were classified into five major functional groups.

**Figure 8.**
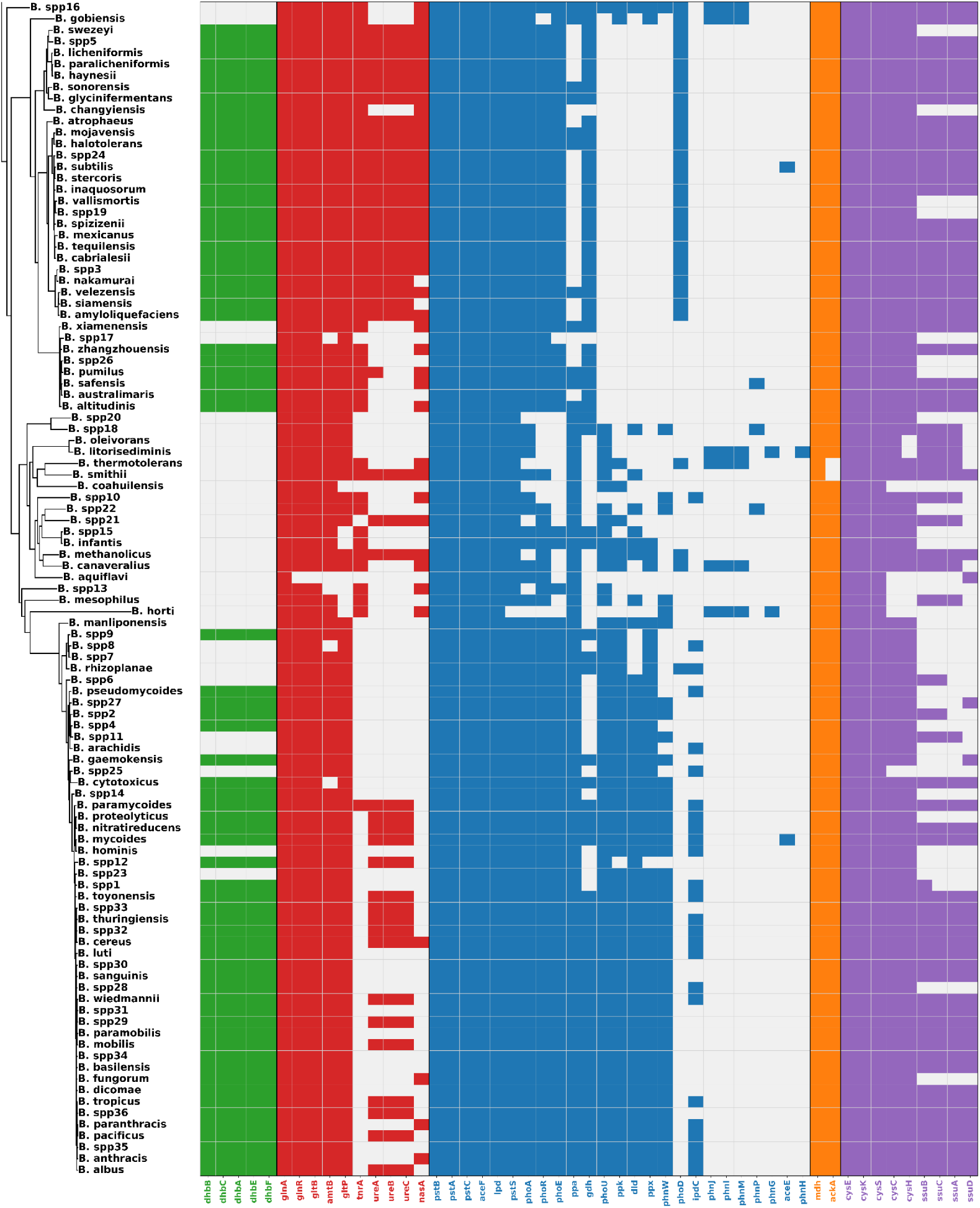
Representative genes associated with biofertilization functions in *Bacillus* species. Genes are grouped into five functional categories: iron acquisition (green, 5 genes), nitrogen acquisition (red, 10), phosphate solubilization (blue, 25 genes), potassium solubilization (orange, 2 genes), and sulfur assimilation (purple, 9 genes). Collectively, these genes underscore the genetic capacity of *Bacillus* to promote plant growth by enhancing nutrient availability, particularly in the rhizosphere.

Iron acquisition was the most broadly represented category and included genes responsible for siderophore-mediated iron chelation, such as those encoding bacillibactin. These genes are critical not only for plant growth promotion but also for antifungal activity, as they deprive pathogens of essential iron (Dimopoulou et al., 2021; Ollinger et al., 2006). Genes in this category were found in nearly all *Bacillus* species analyzed, including *B. licheniformis, B. paralicheniformis, B. subtilis, B. spizizenii*, and *B. velezensis*. However, these genes were absent from genomes of several strains across different *Bacillus* species, including, including *B.* spp20*, B.* spp18*, B. oleivorans, B. litorisediminis, B. thermotolerans, B. smithi, B. coahuilensis, B. spp10, B.* spp22*, B.* spp21*, B.* spp15*, B. infantis, B. methanolicus, B. canaveralius, B. aquiflavi, B.* spp13*, B. mesophilus, B.* spp16*, B. gobiensis, B.* spp17*, B. xiamenensis*, and *B. horti*.

Nitrogen acquisition included key genes for ammonium uptake and regulation (*amtB*, *glnA*, *glnR*, *gltB*), nitrate and nitrite assimilation (*nasA*), urease activity (*ureA*, *ureB*), and associated regulators such as *trnA* (He et al., 2023; Nakano et al., 1995). This potential was observed in most *Bacillus* species, including *B. subtilis, B. stercoris, B. velezensis, B. siamensis*, and B*. spizizenii*, but was less frequent in the *B. cereus* sensu lato clade. Phosphate solubilization encompassed multiple phosphorus acquisition strategies. Acidolysis-related genes (*gdh*, *lpd*, *aceE*, *aceF*, *maeB*, *dlp*), genes encoding phosphatases and polyphosphate-related enzymes (*phoA*, *phoD*, *ppx*, *ppa*, *ppk*, *phnW*, *phnX*). In addition, detected transport and regulatory genes (*phoU*, *phoR*, *phoE pstS*, *pstC*, *pstA*, *pstB*, *phnG*, *phnH*, *phnI*, *phnJ*, *phnM*, *phnP*) are associated with the high-affinity phosphate-specific transport (Pst) and phosphonate (Phn) systems, which facilitate phosphate uptake, recycling, and homeostasis under phosphorus limitation (Pan & Cai, 2023; F. Pang et al., 2024). These genes were nearly universal across species, although phosphorus recycling determinants were generally less abundant.

Potassium solubilization was represented by *mdh* and *ackA*, both involved in potassium mobilization and detected in all species examined (Chen et al., 2022). Finally, sulfur assimilation included genes for sulfur uptake and reduction (e.g., *cysE*, *cysK*, *cysC*, *cysH*, *ssuB*, *ssuC*, *ssuA*, *ssuD*) (Albanesi et al., 2005; Ishikawa et al., 2025), which were also widely distributed across the genus. Overall, these results show that the genetic potential for biofertilization is highly conserved in *Bacillus*, with near-universal representation of phosphate, potassium, and sulfur metabolism genes, while siderophore-mediated iron acquisition and nitrogen assimilation exhibited more variable distributions.

### Promising *Bacillus* lineages for biofungicide development

Between the 103 communities identified, the most promising species were *B. velezensis, B. siamensis, B. amyloliquefaciens, B. spizizenii, B. subtilis*, and *B. atrophaeus*, which showed the highest antifungal potential. These species stand out not only for the presence of genes associated with the production of chitinases, chitosanase, glucanases, and secondary metabolites, but also for lacking virulence determinants in key categories such as exoenzymes, exotoxins, and the T7SS, features that reinforce their biotechnological safety profile.

Regarding antibiotic resistance, *tet(L)* genes were detected in members of the *B. amyloliquefaciens* group; however, their occurrence likely reflects intrinsic genomic features rather than horizontal transfer. A similar pattern may apply to other antibiotic resistance genes, underscoring the need for in vivo studies to evaluate their expression and functional relevance in other agronomically important *Bacillus* lineages, such as *B. pumilus* (Fu et al., 2021)*, B. subtilis*, and *B. licheniformis* (Caulier et al., 2019).

Notably, these promising species have already been experimentally demonstrated to be effective against a wide range of plant pathogens. *B. velezensis* exhibits a broad antifungal spectrum, acting against multiple phytopathogenic genera such as *Aspergillus, Penicillium, Talaromyces, Cryptococcus, Cryphonectria, Helicobasidium*, and *Cylindrocladium*. In addition, lipopeptide compounds produced by *B. velezensis—*for example, bacillomycin D—have shown potent activity against the anthracnose pathogen *C. gloeosporioides*, surpassing the efficacy of conventional chemical fungicides (Devi et al., 2019; Jin et al., 2020; Xu et al., 2016). Similarly, *B. siamensis* has also demonstrated remarkable antifungal efficacy. A strain isolated from the wheat rhizosphere (*B. siamensis* Sh420) produced lipopeptides that strongly inhibited *Fusarium graminearum*, the causative agent of wheat head blight (Hussain et al., 2024). Another isolate (*B. siamensis* AMU03) was effective against *Macrophomina phaseolina*, which causes wilting in various crops, exhibiting antifungal activity through disruption of the fungal cell membrane (Hussain et al., 2024; Hussain & Khan, 2020).

As for *B. amyloliquefaciens* (e.g., strain SQR9), it produces a wide range of antifungal compounds capable of suppressing various soil-borne pathogens. Extracts from *B. amyloliquefaciens* SQR9 inhibited the growth of *Verticillium dahliae, F. oxysporum, F. solan*i, and *Phytophthora parasitica*. These findings indicate that *B. amyloliquefaciens* exerts its biocontrol activity through the production of lipopeptides (iturins, fengicins, surfactins) and siderophores, which interfere with spore germination and compromise the cell integrity of pathogenic fungi (Li et al., 2014).

In addition, *B. spizizenii* (syn: *B. subtilis* subsp. *spizizenii*) has demonstrated significant antifungal effects. For instance, strain BL-59 (identified as *B. subtilis* subsp. *spizizenii*) markedly inhibited mycelial growth of *Colletotrichum sp*. and *Pestalotiopsis sp*. isolated from infected fruits, with reductions of approximately 49.31% and 42.55%, respectively (Duangkaew & Monkhung, 2021). Similarly, *B. subtilis* subsp. *spizizenii* strain MM19 reduced the mycelial growth of *Alternaria alternata,* the causal agent of leaf blight in marigolds, by up to 83% under laboratory conditions (Priyanka et al., 2018). Together, these findings underscore the potential of *B. spizizenii* as a biocontrol agent against phytopathogenic fungi.

*B. subtilis* is one of the most extensively studied species for biocontrol applications, exhibiting broad-spectrum inhibition against a wide range of phytopathogens (Akinsemolu et al., 2024). It has shown activity against *Alternaria solani, F. oxysporum, Rhizoctonia solani*, and *Clarireedia jacksonii*, the causal agent of dollar spot disease (Chen et al., 2023; Kaur et al., 2023; Ramesh et al., 2024; Tuyen et al., 2023). *B. subtilis* produces a variety of antifungal compounds, including lipopeptides and hydrolytic enzymes, that act synergistically to inhibit the growth and development of multiple plant pathogens. Finally, *B. atrophaeus* stands out for its broad antifungal spectrum and stability. Recent studies have shown that *B. atrophaeus* isolates associated with wild apple trees exert strong antagonism against canker-causing fungi, including *Cytospora mali* and *Cryphonectria parasitica* (Bozorov et al., 2024). Moreover, Pisheh et al. (2024) identified iturine A from *B. atrophaeus* strain HNSQJYH170, which displayed potent fungicidal activity against multiple pathogenic fungi, inhibiting more than 80% of the growth of *F. oxysporum, Aspergillus niger, Penicillium chrysogenum,* and *Mucor hiemalis*. Overall, *B. atrophaeus* exhibits a remarkably broad antifungal profile, effective against both soil-borne and and woody-tissue pathogens. These findings highlight the importance of prioritizing certain *Bacillus* species and strains for the development of next-generation biofungicides. In particular, *B. spizizenii* and *B. atrophaeus* emerge as promising new candidates, as species such as *B. velezensis, B. siamensis, B. amyloliquefaciens,* and *B. subtilis* are already established in agricultural applications. The combination of a broad antifungal spectrum, multiple modes of action (including lipopeptides, enzymes, and volatile organic compounds), and genetically safe profiles makes these strains ideal reference models for future biotechnological applications.

## CONCLUSIONS

In conclusion, the integrative framework combining genomic distance metrics (Mash/ANI), network analyses, and label propagation provided a robust strategy for delimiting 103 communities within the genus *Bacillus*, enabling the identification of taxonomically coherent clusters as well as potentially undescribed lineages. Pangenome analysis revealed contrasting patterns of openness and genomic fluidity, with a significant inverse relationship (φ = 0.4499 − 0.4242α). This trend reflects lifestyle-driven adaptive strategies: generalist and opportunistic species maintain open pangenomes with high gene content variability, whereas more specialized taxa exhibit closed pangenomes, reducing genomic flexibility while enhancing niche adaptation. From an applied perspective, groups such as *B. amyloliquefaciens* (Ngalimat et al., 2021), *B. pumilus* (Fu et al., 2021), *B. subtilis*, and *B. licheniformis* (Caulier et al., 2019) harbor extensive repertoires of hydrolytic enzymes, NRPS/PKS clusters, and other secondary metabolite biosynthetic clusters, positioning them as priority candidates for biofertilizer and biocontrol development. Conversely, the coexistence of resistance determinants and virulence factors within the *B. cereus* sensu lato complex underscores the necessity for stringent biosafety assessments and risk evaluation before considering its application in agricultural settings.

## ACKNOWLEDGEMENTS

This work was supported by Fundação Carlos Chagas Filho de Amparo à Pesquisa do Estado do Rio de Janeiro (FAPERJ), Coordenação de Aperfeiçoamento de Pessoal de Nível Superior-Brasil (CAPES; Finance Code 001), Conselho Nacional de Desenvolvimento Científico e Tecnológico (CNPq) and Programa de Apoio à Pesquisa, Inovação e Cultura (PAPIC - UENF). The funding agencies had no role in the design of the study and collection, analysis and interpretation of data and in writing.

