## Supplementary figures for "Integrative phylogenomic and pangenome landscape of *Bacillus*: insights from 10,000 genomes into taxonomy, functional potential, and biotechnological applications"

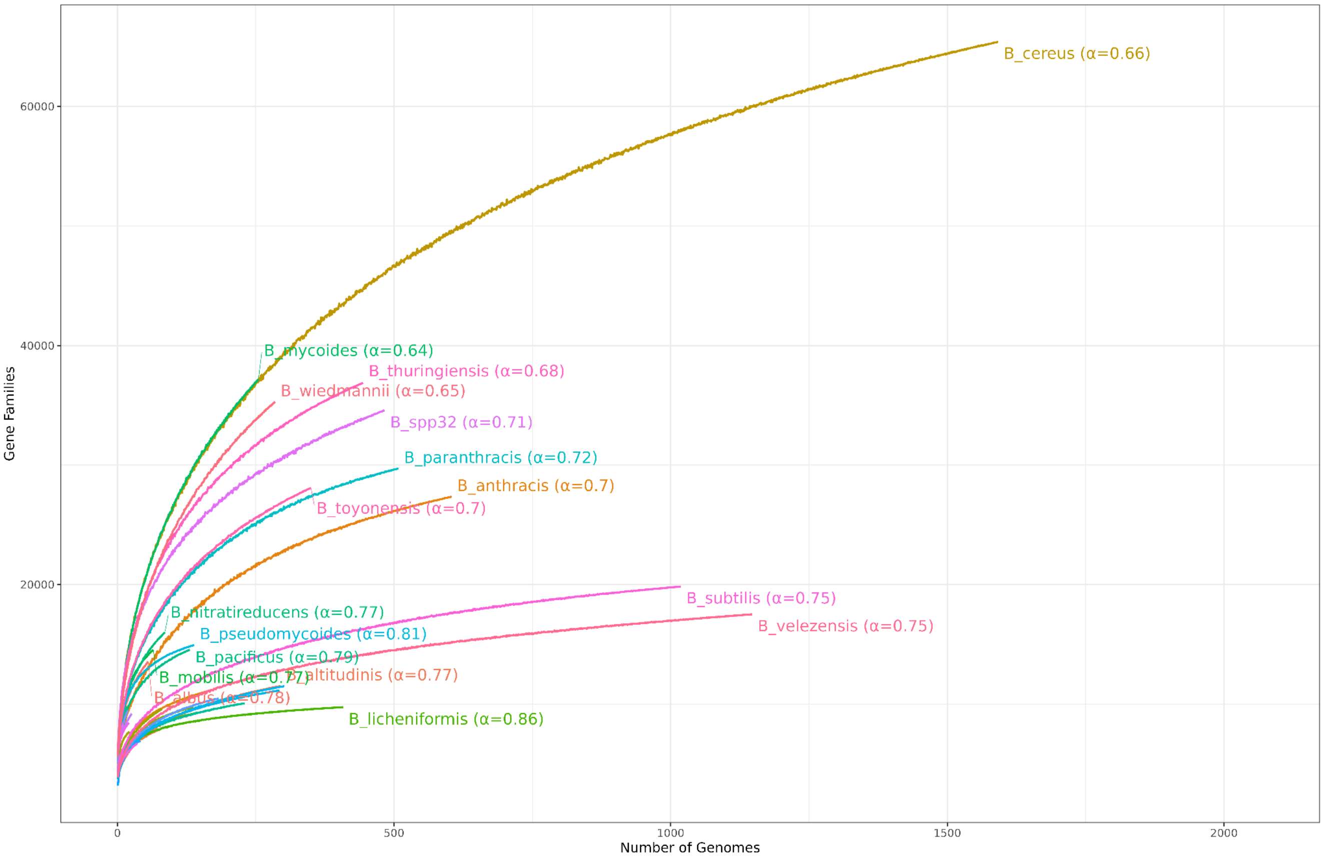


**Figure S1.** Global Bacillus pangenome curves showing gene family accumulation as a function of genome sampling. Each curve represents a distinct Bacillus lineage, with α values indicating the pangenome openness estimated from Heaps' law. Lower α values denote more open pangenomes (greater gene diversity), whereas higher α values indicate more closed pangenomes.

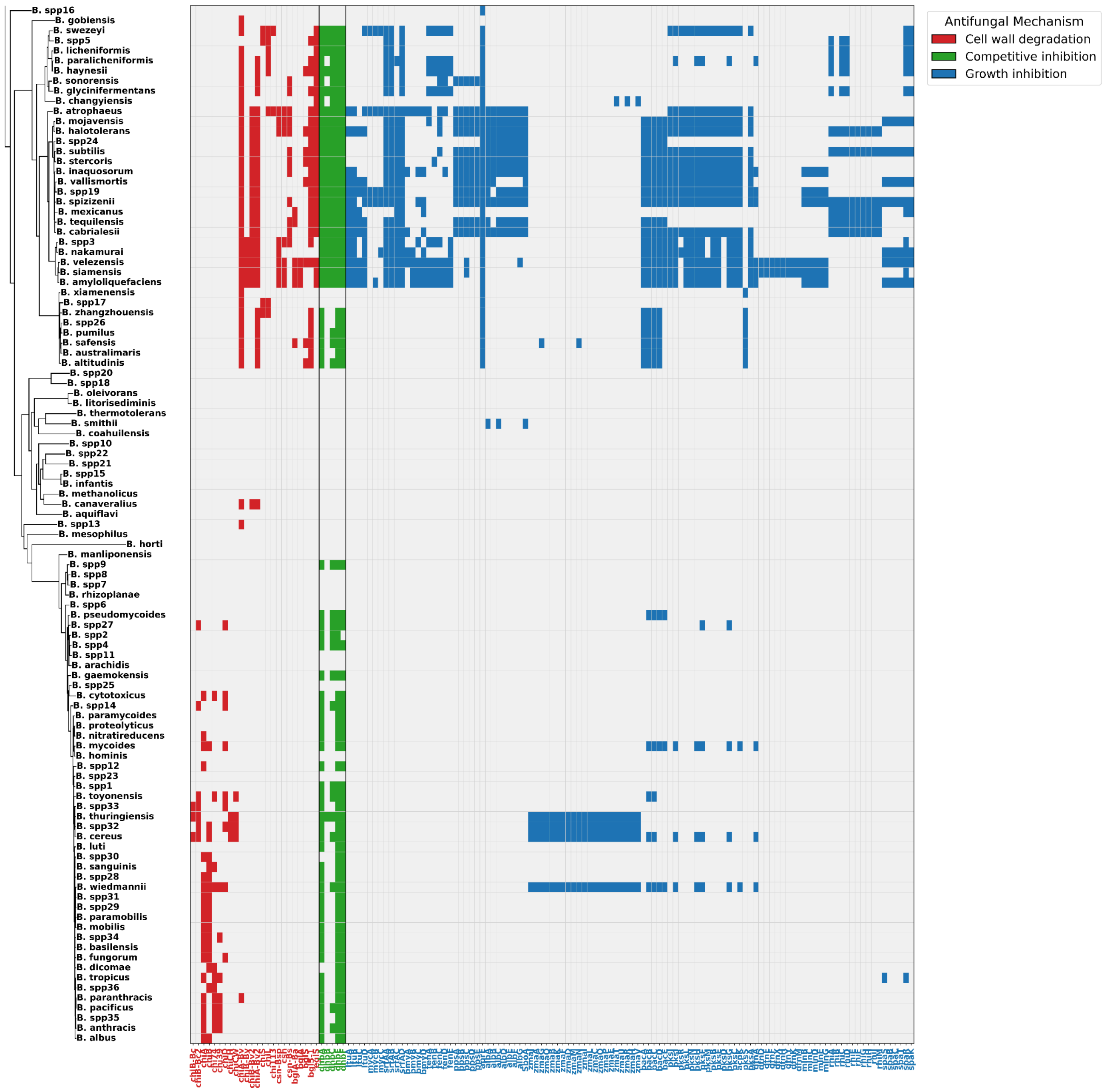


**Figure S2.** Presence-absence matrix of genes with antifungal potential across *Bacillus* identified using USEARCH v11.0.667. Clusters of genes related to secondary metabolite biosynthesis (e.g., *ituA*, *ituB*, *ituC*, *ituD*) and siderophore production (e.g., *dhbA*, *dhbB*, *dhbC*) are highlighted. The analysis also includes lytic enzyme genes such as chitinases (e.g., *chiB-Bc*, *chiL*), chitosanase (e.g., *csn*), and β-glucanases (e.g., *bglA*, *bglS*).
